# Identifying Network Factors Driving Changes in Brain Connectivity During Aging via Stochastic Actor-Oriented Models

**DOI:** 10.64898/2026.10.02.756273

**Authors:** Emma K. Garrison, Roberto C. Sotero, Javier G. Orlandi, the Alzheimer’s Disease Neuroimaging Initiative

## Abstract

Brain network organization emerges from ongoing network dynamics that continuously create, maintain, and reorganize connectivity patterns. At both the structural and functional levels, these dynamics shape network evolution throughout the lifespan. Here, we introduce a data-driven framework based on Stochastic Actor-Oriented Models (SAOMs), which model longitudinal network changes as a process of local edge formation and dissolution driven by local-connectivity factors. Applied to functional and structural brain networks from an aging cohort, the resulting models recovered a set of interacting mechanisms that collectively promote hub formation, short-range connectivity, and modular organization. These factors were derived through a novel data-driven factor-selection procedure rather than specified a priori, allowing network mechanisms underlying longitudinal change to be identified directly from observations. Functional networks exhibited substantially higher rates of change than structural networks, suggesting more frequent opportunities for network maintenance and reorganization. Structural networks additionally showed despecification in the hub-forming mechanisms with cognitive impairment. These results support a view of brain network organization as an actively maintained process emerging from ongoing local connectivity dynamics. More broadly, this framework provides a general approach for studying longitudinal network dynamics through the estimation of network-generating mechanisms over time.

## Introduction

There is a growing body of evidence that brain function emerges from time-varying interactions among distributed regions of the brain rather than from isolated areas acting independently. This represents a key shift in the methodological approaches utilized to study the brain, including a growing interest in network neuroscience methodologies [1– 4]. The field of network neuroscience has revealed consistent patterns of network structure and topology, including robust observations of small-world topologies with high clustering and short path lengths across brain networks [4–6]. These are highly energy-efficient structures, resulting from growing networks constrained by wiring costs [2, 7, 8]. Brain networks are highly modular, with subnetworks constrained by a trade-off of integration vs. segregation [9–11].

Both structural and functional networks are not static and are continuously evolving throughout the lifetime of an individual, with changes in their topologies and structures at various ages [12, 13]. Lifetime network studies, however, have relied mostly on broad trends of network measures rather than on network mechanisms, as the long-term dynamics of the brain are still poorly understood. Many longitudinal studies of network organization often focus on summary measures and then interpret their changes as evidence about underlying processes [14]. This type of study has limitations, as the underlying processes could produce indistinguishable trajectories of summary statistics through competitive or compensatory mechanisms [15]. Indeed, for network observables that might be externally regulated, it is often difficult to distinguish the absence of dynamics from the balance of opposing dynamics, as shown for example in regional activity measures [16].

Regarding these regulatory mechanisms, homeostatic regulation is well-characterized at cellular and circuit scales [17, 18], but frameworks of maintenance and compensation are attributed to differing and incomplete evidence in stable network-level results [19, 20]. Cost considerations point the same way, as brain network topology is metabolically expensive to build [2], and the majority of cortical energy is spent on supporting ongoing baseline activity [21], suggesting that holding a topology in place carries a cost.

Stable network measures do not imply stable networks. Similar values of modularity, efficiency, or clustering can emerge from different patterns of edge rewiring [14]. Hence, understanding brain networks through time, and aging in particular, requires methods capable of separating observed network states from the mechanisms that produce them. To that end, Stochastic Actor-Oriented Models (SAOMs) [22, 23] have been specifically designed for this purpose. SAOM models network evolution as a continuous-time process in which links are sequentially created, retained, or dissolved between observations based on a hypothesized set of network factors. However, traditionally a set of factors must be proposed by the researcher prior to weight estimation. This requires a body of previous research and/or a strong logical local connectivity argument, both of which are absent when considering an application to brain networks. For any known previous attempts to utilize this methodology in neuroscience [24], the approach was still largely inferential, relying on untested hypotheses rather than deriving the factors from the data.

In this work, we introduce a novel data-driven procedure based on SAOM to identify the local-connectivity factors driving change in functional and structural human brain networks during aging. The resulting models point at the underlying processes by which the trends and stability of summary statistics emerge over time, and the parameters can then be compared across demographic and clinical groups, including age, sex, and age-related cognitive impairment, to examine any disruption to this underlying mechanism.

## Methods

### Data Preparation

Data used in the preparation of this article were obtained from the Alzheimer’s Disease Neuroimaging Initiative (ADNI) [database](http://adni.loni.usc.edu). The ADNI was launched in 2003 as a public-private partnership, led by Principal Investigator Michael W. Weiner, MD. The original goal of ADNI was to test whether serial magnetic resonance imaging (MRI), positron emission tomography (PET), other biological markers, and clinical and neuropsychological assessment can be combined to measure the progression of mild cognitive impairment (MCI) and early Alzheimer’s disease (AD). The current goals include validating biomarkers for clinical trials, improving the generalizability of ADNI data by increasing diversity in the participant cohort, and to provide data concerning the diagnosis and progression of Alzheimer’s disease to the scientific community. For up-to-date information, see adni.loni.usc.edu.

All data were selected from the ADNI-3 phase of the study and filtered for participants with scans from more than one time point. There are a total of 260 participants—132 in the control group and 128 with cognitive impairment. The majority of the impaired group have some form of Mild Cognitive Impairment (MCI), and only 9 individuals have Alzheimer’s Disease (AD). 148 participants are female, and 112 participants are male. Participant ages range from 56–92 years of age. The data include both resting-state functional magnetic resonance images (fMRI) to construct functional networks and diffusion tensor images (DTI) to construct structural networks.

Resting-state fMRI data were preprocessed in MATLAB using SPM12 [25], including motion correction, coregistration to each subject’s T1-weighted image, global signal removal [26], nuisance regression, and bandpass filtering (0.01–0.08 Hz). Regional BOLD time series were extracted as gray matter-weighted means within each parcel of the atlas, and Pearson correlation matrices were computed; binary adjacency matrices were derived by retaining the top 20% of connections.

Diffusion MRI data were preprocessed following the MR-trix3 structural connectome pipeline [27], including denoising, eddy current and motion correction, fiber orientation distribution estimation via msmt-CSD [28], and anatomically-constrained probabilistic tractography [29] with SIFT2 streamline weighting [30]. Structural connectomes were constructed using tck2connectome and binary adjacency matrices derived by retaining the top 20% of edges by connection weight. In both cases, the acquisition protocols were kept constant for each participant across all their time points. See supplementary material for additional details.

This study used the Schaefer et al. 2018 functional atlas with 7 subnetworks [10, 31]—Visual Network (VN), Somato-motor Network (SMN), Dorsal Attention Network (DAN), Salience Ventral Attention Network (SN), Limbic Network (LN)^1^, Frontoparietal or Control Network (FPN), and Default Mode Network (DMN). This is a functional atlas but was applied to both types of data for consistency.

The analyses in this study are conducted on the time window level rather than the participant level. A time window is defined as a set of two consecutive scans. There are 421 functional time windows and 329 structural time windows in the dataset. Participants with functional data have an average of 1.6 time windows, and participants with structural data have an average of 1.5 time windows. Repeated measures were considered through mixed-effects linear models [32].

The impairment level of a time window was categorized based on the average Clinical Dementia Rating Sum of Boxes (CDR-SB) [33]. A time window is classified as *control* if the average CDR-SB is less than or equal to 0.5, and as *impaired* if the average CDR-SB is greater than 0.5. Due to the relatively low number of AD participants, the groups were not divided further. There are 225 functional time windows and 175 structural time windows in the control group and 217 functional time windows and 170 structural time windows in the impaired group.

### Stochastic Actor-Oriented Model

This study utilized the Stochastic Actor-Oriented Model (SAOM) as a framework for analyzing the factors driving change on the network. This framework was originally designed to model how social networks change driven by individual choices, and in the context of brain networks, it models how local connectivity drives large-scale changes on the network [22, 23]. This model and its estimations are implemented in the R-based Simulation Investigation for Empirical Analysis (RSiena) [34–36]. The following details describe the model and procedures as implemented in this study.

The SAOM framework simulates changes on a network (***G***) and consists of a rate (*ρ*), a list of factors 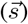, and the weights of those factors 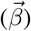. In this study, factors are usually network measures; e.g., the Factor Degree is calculated as the number of nodes to which node *i* is connected. For the full list of factors, see the Factor List subsection below.

In short, the procedure is based on a continuous-time Markov chain and works as follows. For a given time window, at the initial time point, one node (*i*) is randomly selected to have the opportunity to change. The list of possible changes is defined by the objective function (*f*_*i*_),

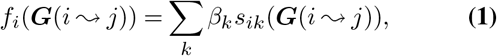

where ***G***(*i*↝*j*) describes the network once the connection between node *i* and node *j* has been changed. If node *i* and node *j* are connected, their connection is dissolved, and if node *i* and node *j* are not connected, a connection is formed between them. *β*_*k*_ is the weight for factor *k*; *β*_*k*_ can be either positive or negative for each factor, indicating a positive or negative priority on the presence of that network structure in the network. *s*_*ik*_ is the factor *k* calculated for node *i*. Note that the objective function also includes the condition of no change occurring. Once the objective function is calculated for every possible change, the probability of making a particular change is calculated by a weighted exponential,

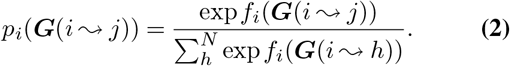

A time step associated to this change, Δ*t*, is selected from an exponential distribution,

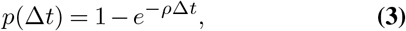

and added to the current time. The procedure is then repeated until the current time reaches the time window length. The rate (*ρ*) and factor weights 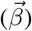 are estimated given the networks at the initial and final time points (*t*_1_ and *t*_2_) through the Generalized Method of Moments and the Robbins-Monro algorithm. This produces an estimated vector 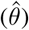 containing *ρ* and 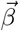. This estimation procedure is implemented in RSiena [36].

To estimate the relevance of a particular factor, we utilize the RSiena standard procedure [34]. In short, the weight of the factor being tested (*k*) is fixed at a particular value (usually 0), and the other factors are estimated. The null hypothesis of these score-type tests is that the weight of the factor k is 0 (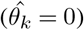), and the statistic is the t-ratio for that factor,

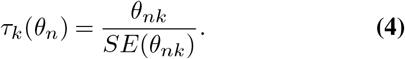

A p-value for this test less than 0.1 suggests that the weight of factor *k* is not zero with a 90% confidence interval. If the hypothesis of the factor weight being zero cannot be rejected, it is likely that the factor can be removed from the overall hypothesis. This test creates the basis for the Factor Selection Procedure.

### Factor Selection

Traditionally, the factors in the SAOM are selected by the researcher before the weights are estimated. This selection is made based on discipline-specific principles, logic, and prior research. We developed a novel data-driven framework specially suited to brain networks in which the factor set does not have to be preselected.

The procedure involves successive factor evaluation scoretype tests. Note that factors in the model are highly interdependent, as they are linked to network measures, so an evaluation of a factor independent of the rest of the hypothesis (the other factors) is not particularly useful. Our procedure was designed to mitigate this interdependence.

#### All-Pairs Initialization

The procedure begins with the *all-pairs initialization* during which each factor is tested with every other factor. For example with three potential factors (*A, B*, and *C*), this would mean 6 tests—*A* tested with *B, A* tested with *C, B* tested with *A, B* tested with *C, C* tested with *A*, and *C* tested with *B*. From these tests, the factor with the lowest average p-value across all its tests, given that *p* < 0.1, is added as the first factor. From there, the factor with the lowest p-value, given that *p* < 0.1, among all the tests with the first factor is added as the second factor. This all-pairs initialization can conclude with 0, 1, or 2 factors. If 2 factors are found, the rest of the procedure continues with the *motion to reconsider* and *factor addition* repeated until the stopping condition is reached.

#### Motion To Reconsider

After a new factor is added to the current hypothesis, a *motion to reconsider* is conducted, testing each factor in the hypothesis to ensure that it still meets the requirement that *p* < 0.1. If the inclusion of a new factor means that the weight of a previous factor in the hypothesis cannot confidently be considered not to be 0, that other factor is removed from the current hypothesis. The factor remains in consideration for future addition to the hypothesis as it evolves.

#### Factor Addition

The next step is to test each of the factors that are not in the current hypothesis for a possible *factor addition*. The factor with the lowest p-value, given that *p* < 0.1, is then added to the hypothesis. This, followed by another motion to reconsider, is repeated until no new factor weight can be considered not to be 0 with a 90% confidence interval. Thus, the current hypothesis contains the minimum set of factors that best explain the data. See supplementary material for additional details.

### Aggregate Factor Selection

The factor selection procedure is conducted on individual time windows; thus, an aggregate set of factors was selected across all time windows for comparisons. Based on model data, an aggregate factor was selected if it appeared in at least 40% of the time windows analyzed.

Following a standard testing/validation procedure, the aggregate factor list was selected from a subset of 40 participants with 53 functional and 59 structural time windows. The remaining 220 participants were the ones used for subsequent analyses. See supplementary material for details on the groups.

The factor selection procedure was validated on a simulated dataset where the sets of factors were known. Validation resulted in an Area Under the Curve for the Receiver Operating Characteristic and Precision-Recall of 0.767 and 0.787, respectively. The procedure generally showed better precision than recall, indicating that it is more likely to miss real factors than to include spurious ones. This is likely due to the correlation between factors and the fact that the procedure was designed to identify the minimum set of factors that best explain the changes. See the supplementary material for full validation analysis.

### Factor List

The total set of factors falls into three categories—Whole Network Factors, Euclidean Distance Factors, and Subnetwork Interaction Factors. The Euclidean distance factors and subnetwork interaction factors are factors from RSiena’s set of covariate factors, with the Euclidean distance indicating the physical distance between brain regions in the atlas and the subnetwork labels being pulled from the atlas’s labels (these values do not change between time points). The factors that were considered in this work included Whole Network Factors: Degree, Transitive Triads, Betweenness, Distance 2, 4-Cycles, and Assortativity; Euclidean Distance Covariate Factors: Euclidean Distance and Euclidean Neighbor Distance; and Subnetwork Covariate Factors: Same Subnetwork, Same Subnetwork Transitive Triplets, Jump Subnetwork Transitive Triplets, Same Subnetwork Distance 2, and Same Subnetwork 4-Cycles. The definitions can be found in the RSiena manual [34] and in the Supplementary Material. In total, thirteen factors were considered. *|*

### Model Evaluation

Once the aggregate set of factors was found for each modality (functional and structural), the weights of those factors were estimated for every time window across the remaining subjects. Hence, each time window resulted in a rate estimation (*ρ*), its standard error, and factor weight estimations 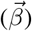 and their standard errors. This data structure led to the implementation of a series of mixed-effects linear models weighted by the inverse variance (1/*SE*^2^) to incorporate the estimation precision, with a random intercept (1 |*ID*) for the subject identity to account for repeated measures. Finally, for Rate analyses specifically, the *Y ears* term is added to the model to account for dependencies in the number of years between the scans, which would impact the Rate. The majority of time windows are 1 year apart, but a few have a longer gap.

Prior to the comparisons, certain time windows were removed due to poor estimation or outlier values. Any window where a factor’s error exceeds 20× that factor’s median error was dropped, as this indicates poor model convergence [35]. Then multivariate outliers were identified using Mahalanobis distance (median/MAD centering, minimum covariance determinant estimation) and excluded if they exceeded the 97.5th percentile of the *χ*^2^ distribution [37]. This removed about 40 time windows from the whole-brain analyses. Due to the different factor sets between the modalities, the factor weights cannot be compared directly. However, the Rate can be compared between the functional models and the structural models using the following mixed-effects linear model:

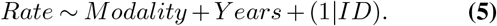

A t-test on the *Modality* coefficient determines difference significance between the functional and structural rates on each of the systems. This can be done because the underlying atlas of nodes is the same.

From there, group differences and age-related trends for the Rate and factor weights were assessed using their own mixed-effects linear models. *Impairment, Sex*, and their interaction were tested as Type-III omnibus terms in one joint model,

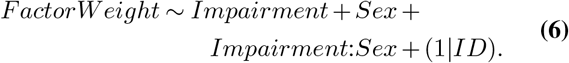

Age trends were tested in a three-way model,

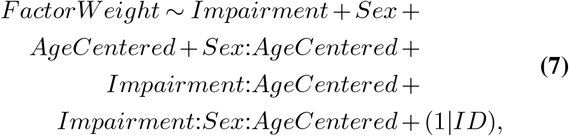

from which the Overall, Control/Impaired, and Male/Female Age Slopes were extracted as marginal contrasts—the Overall slope was averaged over the sample’s actual joint Impairment-and-Sex proportions, and each group-specific slope was evaluated at the sample’s actual proportion of the other factor alone. Interaction terms were defined as the difference between their flanking slopes.

To control for multiple comparisons, p-values were corrected using the Benjamini-Hochberg false discovery rate (FDR) procedure. Correction was applied within families of tests pooled across Rate and all factor weights for a particular comparison.

## Results

Our data-driven framework, developed to characterize longitudinal changes in functional and structural human brain networks during aging, resulted in Factors Degree, Transitive Triads, Betweenness, Distance 2, 4-Cycles, Euclidean Distance, and Euclidean Neighbor Distance as those involved in driving network changes in both functional and structural whole-brain networks. Factors Same Subnetwork and Same Subnetwork 4-Cycles were found only in functional brain networks. Factor Assortativity was found only in structural networks.

Using these factor sets, the remaining time windows found rates for each of the modalities for each of the systems. These rates are reported and compared between the functional and structural in Figure 1.a. The functional rate is significantly higher than the structural rate, indicating that, during a time window, each node in the functional networks has more opportunities ( *∼*70) to change compared to the struc-tural ( *∼*20).

**Fig. 1.**
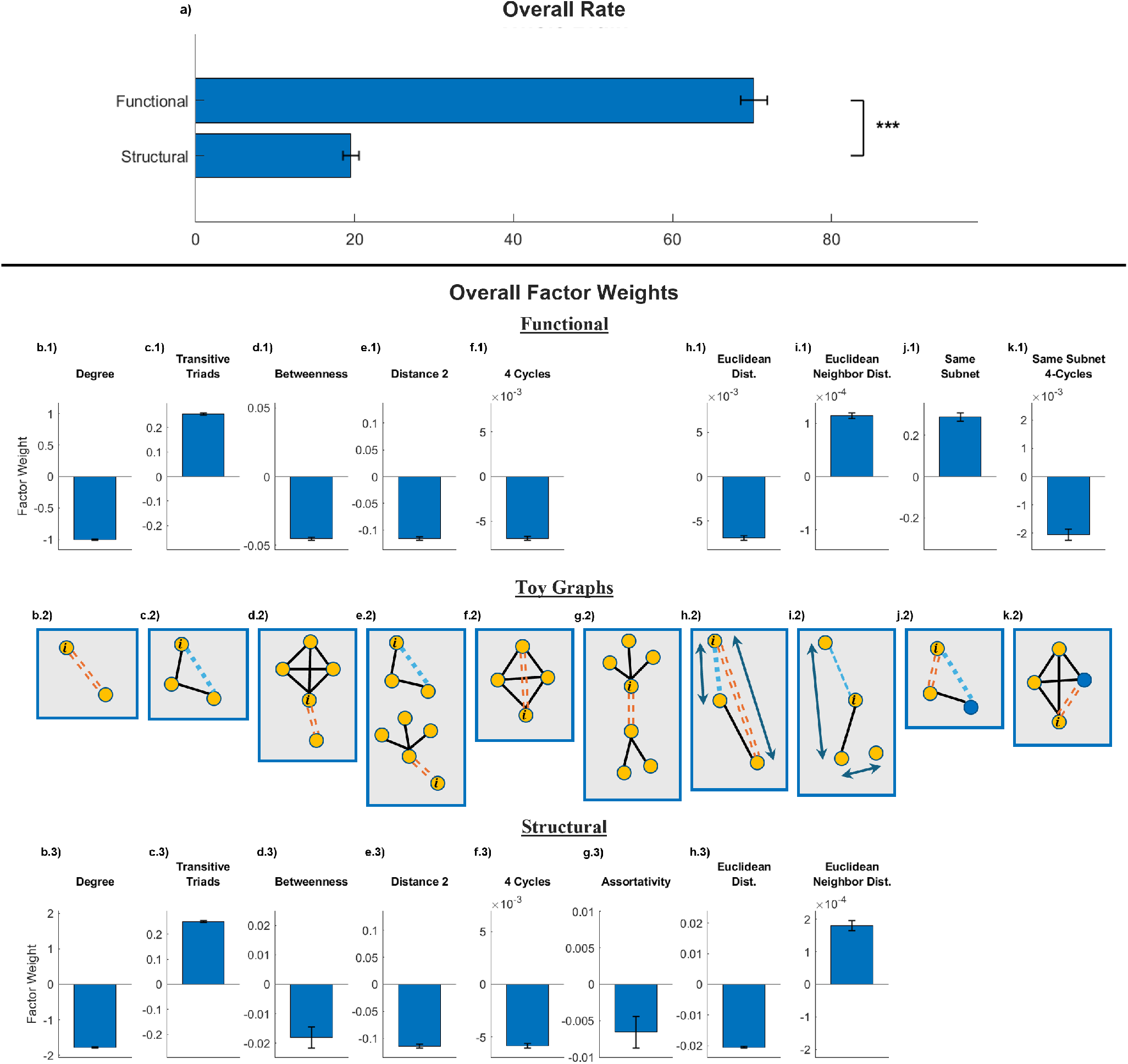
Overall Results. Overall results for the rates and factor weights. The rates are compared between structural and functional (a), combining all examples for a mixed-effects model controlling for years between scans and including subject ID as a random intercept. There is a significant difference between the functional and structural rates (*p* = 6.2 × 10^−118^). Factor weight results for the whole-brain models for the functional (b-f.1 and h-k.1) and the structural (b-i.3). Bar graphs represent *µ ± σ* values for the factors computed via two-stage inverse-variance weighting. Factors with a positive weight are positively prioritized, and factors with a negative weight are negatively prioritized. Between the functional and structural are Toy Graphs (b-k.2) that demonstrate the local expected behaviors based on the signs of the factors. A blue single dashed line indicates a likelihood to build a connection, while an orange dashed double line indicates a likelihood to dissolve a connection.

Functional network factor weights are shown in Figure 1 (b-f.1 and h-k.1) and structural ones in Figure 1 (b-i.3). The sign of factors was consistent across modalities. In particular, we observed a negative weight for Factor Degree, indicating nodes are prioritizing not making new connections (Figure 1.b). A positive weight for Factor Transitive Triads suggests that nodes are prioritizing closing open triangles (Figure 1.c). A negative weight for Factor Betweenness means that nodes are prioritizing disconnecting from nodes that are not connected to their neighbors (Figure 1.d). A negative weight for Factor Distance 2 implies that nodes are building connections to close open triangles or, more often, disconnecting from nodes that are connected to many nodes to which the node is disconnected (Figure 1.e). A negative weight for Factor 4-Cycles indicates that nodes are prioritizing disconnecting from redundant connections inside of 4-cycles (Figure 1.f). A negative weight for Factor Assortativity, which is only present in the structural models, indicates that high-degree nodes are prioritizing disconnecting from other high-degree nodes (Figure 1.g). A negative weight for the Factor Euclidean Distance shows that nodes are prioritizing connecting to physically close nodes or disconnecting from physically distant nodes (Figure 1.h). A positive weight for Factor Euclidean Neighbor Distance means that nodes are prioritizing connecting with nodes that are physically far from their network neighbors (Figure 1.i). A positive weight for the Factor Same Subnetwork implies that nodes are prioritizing connecting with nodes in the same subnetwork and disconnecting from nodes in different subnetworks (Figure 1.j).

And finally, a negative weight for Factor Same Subnetworks 4-Cycles suggests that nodes are prioritizing disconnecting from redundant connections that pass through nodes of a different subnetwork (Figure 1.k).

These factors act together and sometimes compete for the highest priority. Note that the weight values for each of the factors are on different scales from one another and cannot be compared directly.

One of the central motivations for using SAOMs is that similar network states can arise from different underlying processes. Consequently, after identifying the factors that best explain longitudinal network evolution, we asked whether those factors are equally prioritized across demographic and clinical groups. Differences in factor weights would suggest distinct network adaptation or maintenance strategies despite potentially similar network-level outcomes. We therefore evaluated age-, sex-, and impairment-related differences in both Rate and factor weights using mixed-effects models as described in the Methods section.

The resulting demographic effects are summarized using the t-weighted Effects Direction metric and summarized in Figure 2 and Figure 3 for functional and structural brain networks, respectively. The t-weighted Effects Direction metric indicates whether a certain factor is moving away (positive, purple) or towards (negative, orange) 0 in that particular group. In other words, this graph indicates whether the factor is being prioritized more or less in that group regardless of whether that prioritization is positive or negative.

**Fig. 2.**
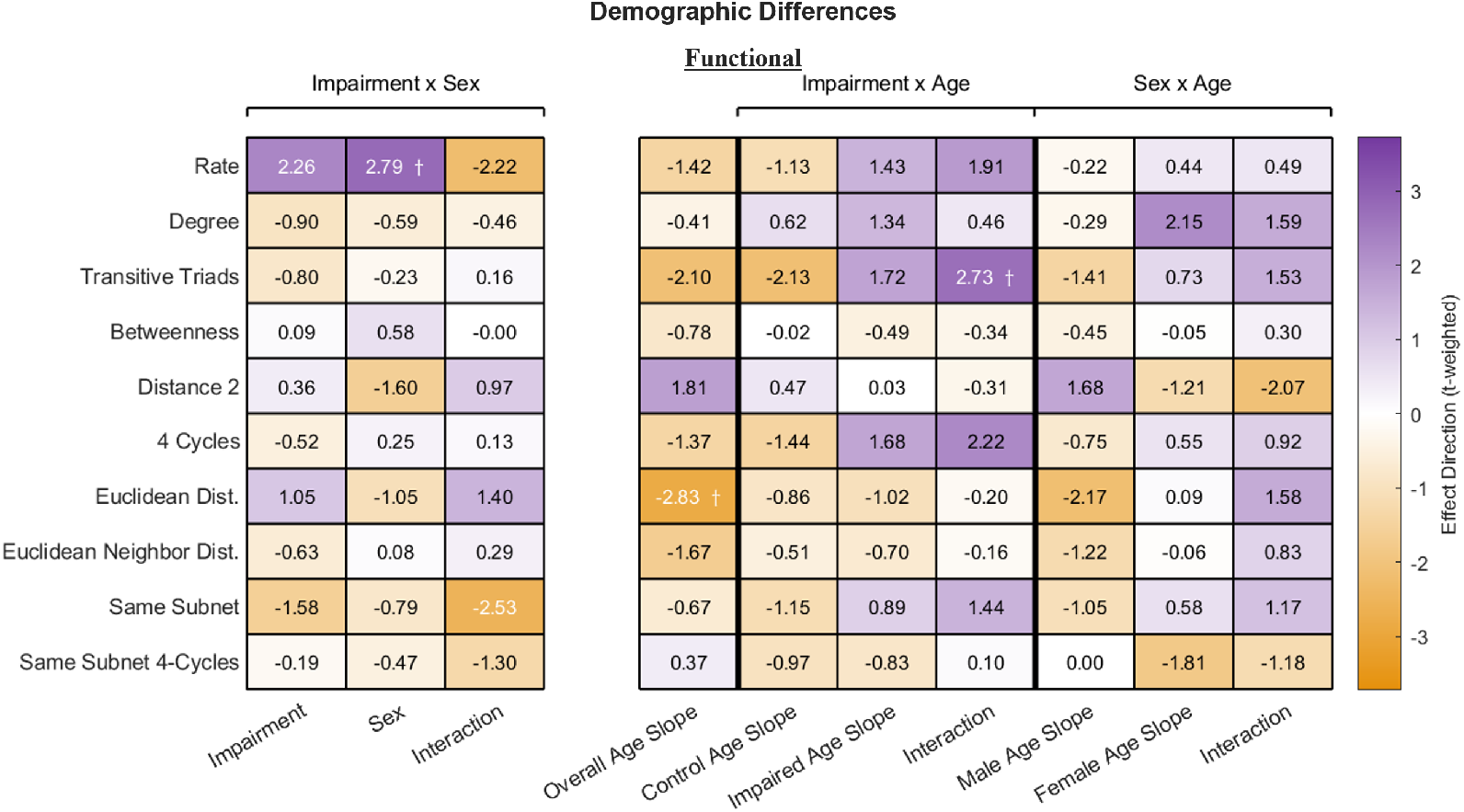
Functional Demographic Results. Results from mixed-effects models comparing whole-brain functional Rate and factor weights across *Age, Sex* (Female–Male), and *Impairment* (Impaired–Control). Effect direction reflects increased (positive) or decreased (negative) prioritization, independent of the factor weight’s sign. Columns 1–3 are Type-III omnibus terms from one joint model of *Sex, Impairment* and their interaction. Columns 4–10 are marginal slopes and their exact difference, all from one joint model. The Overall slope is marginalized over the sample’s actual joint Impairment-and-Sex proportions; each group-specific slope is marginalized over the sample’s actual proportion of the other factor alone. Rate models additionally include a years-between-scans covariate (+*Y ears*). Three results reached marginal significance after FDR correction: a trend toward higher whole-brain functional Rate in females (*p* = 0.056), an *Impairment × Age* interaction for Transitive Triads weight (*p* = 0.068), and the overall age slope for Euclidean Distance has a p-value that rounds to 0.1 (*p* = 0.099), meaning that it is not a real marginal result. *†* represents *p <* 0.1, *\** represents *p <* 0.05, *\*\** represents *p <* 0.01, and *\* * \** represents *p <* 0.001.

**Fig. 3.**
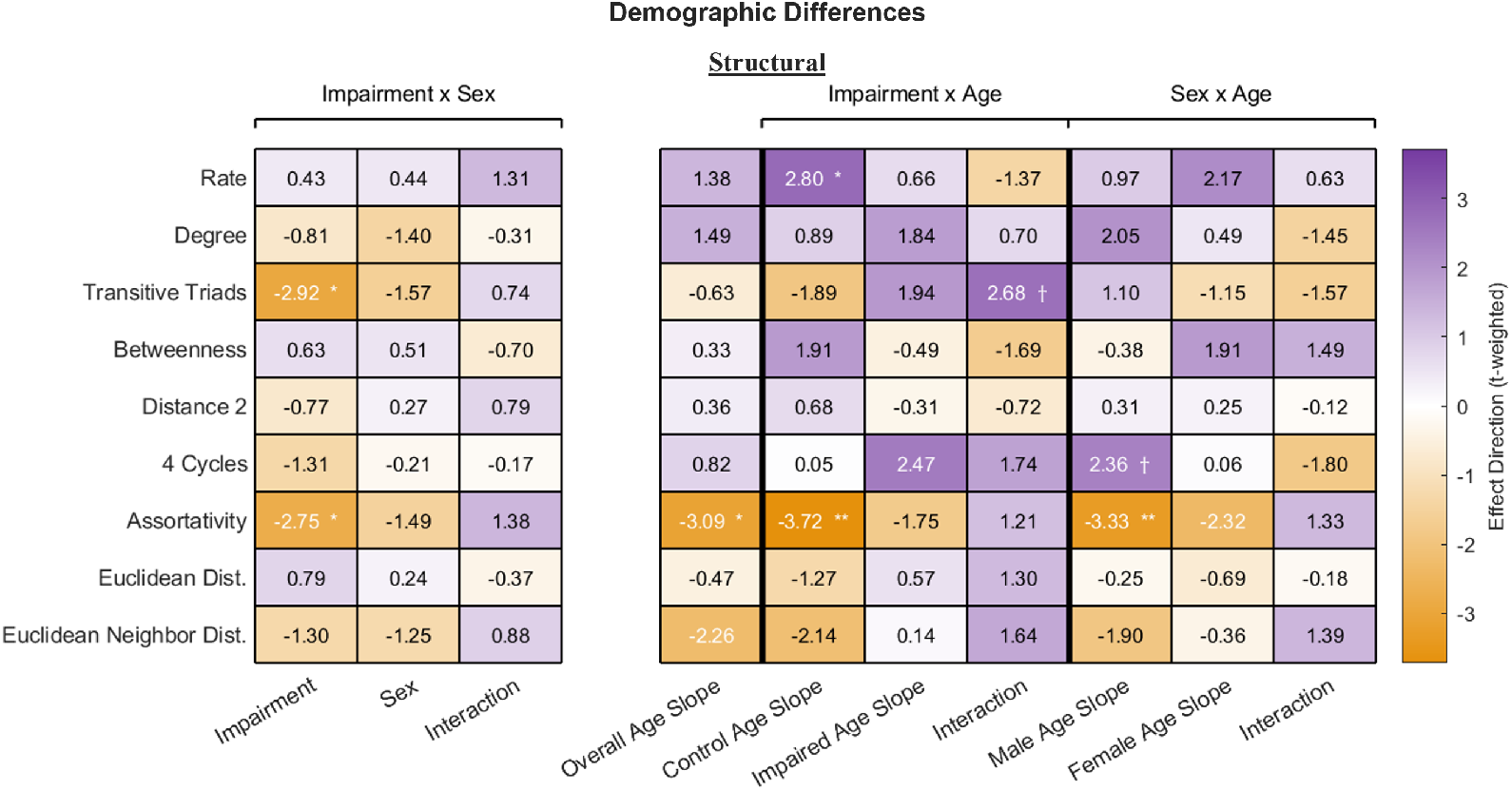
Structural Demographic Results. Results from mixed-effects models comparing whole-brain structural Rate and factor weights across *Age, Sex* (Female–Male), and *Impairment* (Impaired–Control). Effect direction reflects increased (positive) or decreased (negative) prioritization, independent of the factor weight’s sign. Columns 1–3 are Type-III omnibus terms from one joint model of *Sex, Impairment* and their interaction. Columns 4–10 are marginal slopes and their exact difference (interaction), all from one joint model. The Overall slope is marginalized over the sample’s actual joint Impairment-and-Sex proportions; each group-specific slope is marginalized over the sample’s actual proportion of the other factor alone. Rate models additionally include a years-between-scans covariate (+*Y ears*). Six results reached significance after Benjamini-Hochberg FDR correction: a higher rate at higher ages in the control group (*p* = 0.025), a lower weight for Transitive Triads in the impaired group (*p* = 0.029), a less negative weight for Assortativity in the impaired group (*p* = 0.029), a less negative weight for Assortativity at higher ages overall (*p* = 0.021), in the control group (*p* = 0.0024), and among males (*p* = 0.0092). Two results reached marginal significance after FDR correction: *Impairment × Age* interaction for the weight of Transitive Triads (*p* = 0.072) and a more negative weight for 4-Cycles at higher ages among males (*p* = 0.087). *†* represents *p <* 0.1, *\** represents *p <* 0.05, *\*\** represents *p <* 0.01, and *\* * \** represents *p <* 0.001.

The functional models only showed marginal differences on certain measures with no significant trends after FDR correction (Figure 2). Specifically, females exhibited a trend toward higher Rate than males, suggesting a modest increase in the number of opportunities for network reconfiguration over a given time interval. We also observed a marginal *Impairment ×Age* interaction for the weight of Factor Transitive Triads, indicating that age-related changes in triadic closure may differ between cognitively impaired and control participants. Taken together, these findings suggest that the mechanisms driving longitudinal functional network evolution are largely conserved across demographic and clinical groups, with only limited evidence for systematic differences in factor prioritization.

In contrast, the structural models revealed several significant demographic effects (Figure 3). First, the Rate increased with age in the control group, indicating that structural networks undergo more rapid reorganization during healthy aging. Second, the weight of Factor Transitive Triads was lower in the impaired group compared to the control group, suggesting a reduced prioritization of local triadic closure in structurally evolving networks. Third, Factor Assortativity had a less negative weight in the impaired group, implying weaker discouragement of hub-to-hub connectivity. A similar reduction in the weight of Factor Assortativity was observed with increasing age across the full sample, within the control group specifically, and among males. These results suggest that aging and cognitive impairment are associated with changes in the mechanisms governing structural network evolution, particularly those regulating local clustering and hub connectivity.

## Discussion

The models estimated here describe network change as a process rather than as an outcome. Through the SAOM framework, network evolution is represented as a sequence of local edge modifications occurring between observations, allowing the mechanisms governing change to be estimated directly. Each model specifies a set of local connectivity factors that together determine the conditions under which a given connection is more or less likely to form or dissolve. These factors are not summaries of how the networks differed across time, but rather estimates of the mechanisms governing network changes. The derived models therefore report what drove change rather than what changed. A factor’s prioritization therefore describes a pressure acting on the network rather than a change in an observable network property. The factors often oppose and interact with one another, and the effect on any observable property depends on the full set in combination.

Specifying this model has previously required committing in advance to which mechanisms matter, with the inclusion or omission of factors predetermining their possible interactions. Such a predetermination was not made here, and the resulting model from our data-driven procedure contained factors consistent with those established independently in the literature—small-world topology [4–6], wiring efficiency [2, 7, 8], and modularity [9–11], assembled from the local connectivity rules these models derived. The results were consistent across participants and interpretable in the context of brain connectivity.

The interpretive frame follows from factor definitions. In the absence of any factors, these networks would trend toward maximally random graphs; each factor here is a constraint that biases the network away from randomness and toward some particular organization. The derived factors can therefore be read as the constraints under which network structure is held, and their combination as a description of what is being held, consistent with maximum entropy principles.

Several constraints govern global properties. Because all networks are thresholded at 20% density, factors discouraging connection should outnumber those encouraging it, and they do. Factor Degree discourages connection generally and is likely a principal driver keeping density near threshold. Factor Transitive Triads encourages triad closure and, over time, hub formation—the basis of the short path lengths characteristic of small-world topology [38]. Factor Cycle-4 discourages redundancy in square motifs, limiting the clique sizes that Factor Transitive Triads would otherwise produce and thereby moderating subnetwork size.

A second group of factors governs segregation. Factors Betweenness and Distance 2 both favor network partitioning, and together with Factor Transitive Triads, provide the local mechanisms necessary for modular organization: dense connectivity within communities and sparse connectivity between them [39]. In the functional models, Factors Same Subnetwork and Same Subnetwork 4-Cycles further constrain these processes to align with canonical subnetworks. The structural models instead contain Assortativity, which discourages hub-to-hub connectivity and appears to provide a complementary constraint on hub growth in the absence of subnetwork-specific factors.

A third group of factors governs spatial cost. Factor Euclidean Distance favors short-range connections, while Factor Euclidean Neighbor Distance promotes connections that extend beyond a node’s immediate spatial neighborhood. Together they describe a network that balances wiring economy with global reach. This result is consistent with established evidence that long-range connections are metabolically and materially expensive, particularly in structural networks [2, 21]. More importantly, the models describe these spatial constraints as ongoing pressures acting on network evolution rather than simply properties of the final network configuration. Sustaining network organization may therefore require the continual suppression of costly longrange rewiring, extending wiring-cost arguments from network structure to network maintenance.

In summary, these factors describe a network maintained under simultaneous constraints of density, modularity, and wiring cost. The resulting organization consists of segregated communities with highly connected hubs, and a spatial layout that balances topological efficiency against wiring cost.

The functional and structural models differed substantially in Rate, that is, the number of opportunities the network has to change within a given period (Figure 1.a), with the functional networks showing a much higher rate. This by itself is not surprising, as one would expect functional changes to occur at a much faster rate or timescale than structural ones. Read as part of a maintenance mechanism, however, this difference could be interpreted as a kind of correctional budget—how often the system can act to return a perturbed network toward its preferred organization or prevent reversion towards random structure. These functional changes also seem to go hand in hand with compensatory mechanisms, as these networks differed less across demographics and clinical groups.

It is also worth noting that the changes described here focus mainly on the strongest weights. Since the underlying networks are weighted, the building and dissolving of connections reflects movement of edge weights across the chosen 20% threshold rather than the creation or elimination of connections outright. The high rates in the functional models are therefore better understood as fluctuations near threshold rather than as wholesale restructuring, where recently dissolved connections could be sitting just below the boundary rather than completely disappearing.

Regarding structural models, they showed reduced prioritization of two factors in the impaired group relative to controls—Transitive Triads and Assortativity (Figure 3). These factors ordinarily operate in tension, with Factor Transitive Triads promoting the formation of highly connected hubs and Factor Assortativity limiting hub-to-hub connectivity. Their joint reduction does not shift the balance between them, but instead relaxes the constraint that the balance imposes. What is lost is the specification itself. Literature shows some evidence of altered hub connectivity on the structural network with Mild Cognitive Impairment, but reports diverge on whether the hub core is impacted [40] or if feeder and peripheral connections degrade [41], a disagreement that is challenging to adjudicate due to rich-club calculations depending on choice of null model and normalization [42]. Our result describes hub behavior de-specification without relying on rich-club normalization, so it is not subject to these specific sensitivities. A reduction in hub-to-hub discouragement without the decline in the weight for Factor Transitive Triads appears at higher ages within the control group, suggesting that older adults are less likely to encourage dissolving connections between hub nodes. This is consistent with the finding in the literature that rich-club organization is largely preserved in healthy aging [43, 44] even if other measures like global efficiency decline [43]. In the control group, the hub-building pressure remains intact while its counterweight relaxes, so hub organization is maintained or reinforced; in the impaired group, both are reduced regardless of age, so hubs are much less governed altogether. These results in the model do suggest that Mild Cognitive Impairment involves unique mechanisms and is not equivalent to accelerated aging.

Several limitations follow from the modeling framework. The most consequential is that the factor selection procedure arbitrates among a predefined set of candidate factors. It can determine which specified mechanisms are prioritized, but cannot recover a mechanism that was never a candidate. The correspondence between the recovered factors and known network organization is therefore evidence that the procedure selects meaningfully among plausible mechanisms, not that the selected set is complete. The SAOM also assigns all nodes a common set of priorities, adjustable only through covariate effects. The dataset further constrains the scope of the conclusions, as all participants were older adults and imaging data were acquired under multiple scan protocols that were not explicitly modeled.

The analyses also treat time windows cross-sectionally, with repeated measures handled statistically rather than modeled directly. A fully longitudinal SAOM formulation incorporating multiple scans per participant would more closely match the underlying process being estimated. Likewise, the comparison of functional and structural Rates was limited to group-level differences between modalities. Future work could extend the framework to jointly model functional and structural networks, enabling direct tests of the proposed correctional budget hypothesis and of functional-structural interactions more broadly. More generally, the framework is readily applicable to other populations, developmental stages, and progressive conditions, where its ability to derive candidate mechanisms from data rather than specify them a priori may prove particularly valuable.

This study introduces a data-driven SAOM framework for characterizing long-term evolution in functional and structural brain networks. Rather than specifying candidate mechanisms in advance, the framework derives the factors governing network change directly from longitudinal data. When applied to an aging cohort, the recovered factors correspond to established principles of brain organization, including modularity, small-world topology, and wiring-cost constraints. The resulting models provide a mechanistic description of brain network evolution, while the particular factors and interactions identified here characterize the dynamics of the aging population under study.

## Ethics

This work did not require ethical approval from a human subject or animal welfare committee.

## Data Accessibility

Data used in preparation of this article were obtained from the Alzheimer’s Disease Neuroimaging Initiative (ADNI) database (adni.loni.usc.edu). Code used to preprocess the data is available at https://github.com/emma-garrison/MRIPreprocessingPipeline. Code used to implement the SAOM model and generate the results is available at https://github.com/emma-garrison/FactorSelectionProcedure.

## Declaration of AI Use

AI was used to refine and organize code to improve readability and structure; all code and analyses were designed and implemented by the researchers and no data was shared with AI models. AI was used to improve readability and language in the manuscript.

## Authors’ Contributions

E.G.: conceptualization, data curation, formal analysis, investigation, methodology, software, validation, visualization, writing—original draft, writing—review and editing; R.S.: conceptualization, funding acquisition, project administration, resources, software, supervision, validation, writing— review and editing; J.O.: conceptualization, funding acquisition, project administration, resources, software, supervision, validation, writing—review and editing.

## Footnotes

1 This subnetwork was ignored for these analyses as it only contains 5 nodes.

