## Supplementary Material for "Identifying Network Factors Driving Changes in Brain Connectivity During Aging via Stochastic Actor-Oriented Models"

### Supplementary Note S1: Data Preprocessing

**A fMRI Preprocessing** **Volume preparation.** 4D functional images were split into 3D volumes, with the first 10 volumes discarded to allow for signal stabilization.
**Motion correction.** Motion correction was performed using SPM *realign*. The six motion parameters and their squares were saved for subsequent nuisance regression.
**Slice timing correction.** Slice timing correction was omitted given the mixed acquisition protocols present in the dataset with differing TR values ($TR = 3$ s and $TR = 0.607$ s). Acquisition protocol was held constant within each subject across all longitudinal timepoints.
**Coregistration and reslicing.** Functional images were coregistered to the nearest temporally-matched T1-weighted structural image using a normalized mutual information cost function (SPM *coreg*) and resliced to 2 mm isotropic voxels.
**Global signal detrending.** Global signal removal was applied to the resliced time series using the Linear Model of Global Signal (LMGS) method (*cspm_lmgs*) [1].
**Nuisance regression.** Motion parameters (6 parameters and their squares) and mean white matter and CSF signals were regressed from the time series. White matter and CSF signals were derived from binary tissue masks generated during SPM12 *segmentation* of each subject's T1-weighted image.
**Regional signal extraction.** Regional BOLD time series were extracted as gray matter probability-weighted means within each atlas parcel, using the continuous gray matter tissue probability map from SPM12 *segmentation* as voxel weights. Voxels returning NaN values following coregistration were excluded from the regional mean on a per-voxel basis.
**Bandpass filtering.** Bandpass filtering of 0.01-0.08 Hz was applied to the regional time series.
**Spatial normalization and smoothing.** Spatial normalization and smoothing were omitted to preserve native-space anatomical specificity.
**Connectivity and thresholding.** Pearson correlation matrices were computed across all parcel pairs. A proportional density threshold of 20% was applied, retaining the top 20% of connections by absolute weight; under this threshold, all retained connections were positive, and no additional sign-based filtering was applied.

**B DTI Preprocessing** Acquisition protocol (single-shell or multiband multi-shell) was held constant within each subject across time points. **Denoising.** Thermal noise removal was performed using the MP-PCA algorithm (*dwidenoise*) [2].
**Gibbs ringing removal.** Gibbs ringing artifacts were removed using *mrdegibbs* [3].
**Motion and eddy current correction.** Motion and eddy current correction were performed using FSL *eddy* [4] via the MRtrix3 *dwifslpreproc* wrapper, with the *–slm=linear* and *–data_is_shelled* flags specified. Reversed phase-encoding susceptibility distortion correction was performed using FSL *TOPUP* [5], where $b_0$ volumes with opposing phase-encoding directions were present in the scan header (*–rpe_header*). For acquisitions where this information was unavailable, correction proceeded without susceptibility distortion correction.
**Bias field correction.** Bias field correction was applied using the ANTs N4 algorithm (*dwibiascorrect ants*) [6].
**Brain masking.** A brain mask was generated using *dwi2mask*.
**Fiber orientation estimation.** Tissue response functions were estimated using the Dhollander algorithm [7]. Fiber orientation

* Data used in preparation of this article were obtained from the Alzheimer's Disease Neuroimaging Initiative (ADNI) database (adni.loni.usc.edu). As such, the investigators within the ADNI contributed to the design and implementation of ADNI and/or provided data but did not participate in analysis or writing of this report. A complete listing of ADNI investigators can be found here.

distributions were estimated using multi-tissue constrained spherical deconvolution (*msmt-CSD*) [8].
**Intensity normalization.** Global intensity normalization across tissue compartments was performed using *mtnormalise* [9].
**Tissue segmentation and coregistration.** Five-tissue-type segmentation of each subject's T1-weighted image was performed using FSL *FAST* via *5ttgen* [10]. A mean $b_0$ image was extracted from the preprocessed diffusion data and registered to the T1 image using *FLIRT* with 6 degrees of freedom [11]. The resulting transformation was used to bring the five-tissue-type image into diffusion space.
**Tractography.** Anatomically-constrained probabilistic tractography (*ACT*) [10] was performed using the *iFOD2* algorithm, seeded from the gray matter–white matter interface (*gmwmi*), generating 10 million streamlines with backtracking enabled. Maximum streamline length was set to 250 $\mathrm{mm}$ with an FOD amplitude cutoff of 0.06.
**Streamline filtering.** Streamline weights were assigned using *SIFT2* [12] applied to the full 10 million streamline set. Streamlines were additionally filtered to 200,000 using *tckedit*.
**Connectome construction.** Structural connectomes were constructed using *tck2connectome*, with edges symmetric, zero-diagonal, scaled by inverse node volume, and weighted by *SIFT2* streamline coefficients. The atlas parcellation was converted to diffusion space prior to connectome generation.
**Thresholding.** A proportional density threshold of 20% was applied to the weighted connectome, retaining the top 20% of edges by connection weight, and the resulting adjacency matrix was binarized.

### Supplementary Note S2: Factor Selection Procedure

For increased clarity of the Factor Selection Procedure, Figure S1 shows an example of how the procedure might behave with a set of five potential factors. The table on the left represents the tests that are conducted, with the top (dark red) row being the tested factors. Each of the rows below represents an individual set of tests being run on that column's factor. For example, in Frame 1 (all-pairs initialization), Factor $A$ is being tested in a hypothesis with Factor $B$, another test with Factor $C$, and so on down the column. The bottom row is the average p-value from each of the tests in that column; this average is only relevant during the all-pairs initialization. The boxes on the right represent the various factors, and the blue line encircles the hypothesis at the end of the tests conducted in that frame. For example, in Frame 1 the lowest average p-value is for Factor $D$ as seen in the bottom row of the table, so Factor $D$ is in the current hypothesis. One can see that during the second Motion to Reconsider, Factor $D$, which had previously been added, could no longer be considered not to be zero ($p > 0.1$) given that the hypothesis now contains Factor $B$ and Factor $E$. As a result, Factor $D$ is reconsidered and removed from the current hypothesis. The procedure ends when no new factor meets the criterion that $p < 0.1$ (Panel 6).

#### A Thresholding and Estimation

***A.1. Factor Group Selection*** The dataset was split into two groups —the Factor Group for determining the aggregate set of factors and the Estimation Group for estimating and comparing the weights of the factors. These were both tested to make sure that they were representative of the whole group's distribution of impairment, sex, and age.
The Factor Selection Procedure was completed on a subset of the time windows in the dataset. In order to select the most representative sample of those with a completed procedure for the Factor Group, an algorithm that greedily assembles a proposed Factor Group subject set one subject at a time. At each step, it randomly chooses among the top-scoring eligible candidates, where the score balances how few uncomputed windows the subject had and how much they'd help balance the sex and impairment distribution of the current set. Once the sample hits the minimum number of windows (about 12% of the total number of functional or structural windows), it runs $\chi^2$ tests of independence (on sex and impairment) and a Kolmogorov-Smirnov test (on Age) comparing the proposed Factor Group and the proposed Estimation Group to the population. If any test fails ($p < 0.05$), it keeps growing the sample and rechecks. This is repeated many times, and about 500 unique sets were compared on their uncomputed windows and their similarity to the population on sex, impairment, and age, selecting the most representative sample. The chosen Factor Group and Estimation Group were considered fairly representative, with all p-values greater than 0.3. The Factor Group compared to the population on sex with $p = 0.42$, on impairment with $p = 0.49$, and on age with $p = 0.31$. The Estimation Group compared to the population on sex with $p = 0.63$, on impairment with $p = 0.55$, and on age with $p = 1.00$.

***A.2. Thresholding Comparisons*** The Factor Group results for the factors were also tested to determine if there was any significant difference between demographic groups on which factors would be included and excluded. Those results for the whole brain are shown in Figure S2. Yates's corrected $\chi^2$ tests of the proportion of the factor found in each group was conducted for every factor for control/impairment groups, male/female groups, and younger/older groups (cut off a 75 years). All results with $p < 0.1$ are shown, but few are relevant in that they would tip the difference of inclusion or exclusion above or below the threshold. The only relevant significant ($p < 0.05$) difference was found in the inclusion or exclusion of the Factor Same Subnetwork in the structural models in different age groups. These tests were not corrected for repeated measures or interaction between impairment, sex, and age.

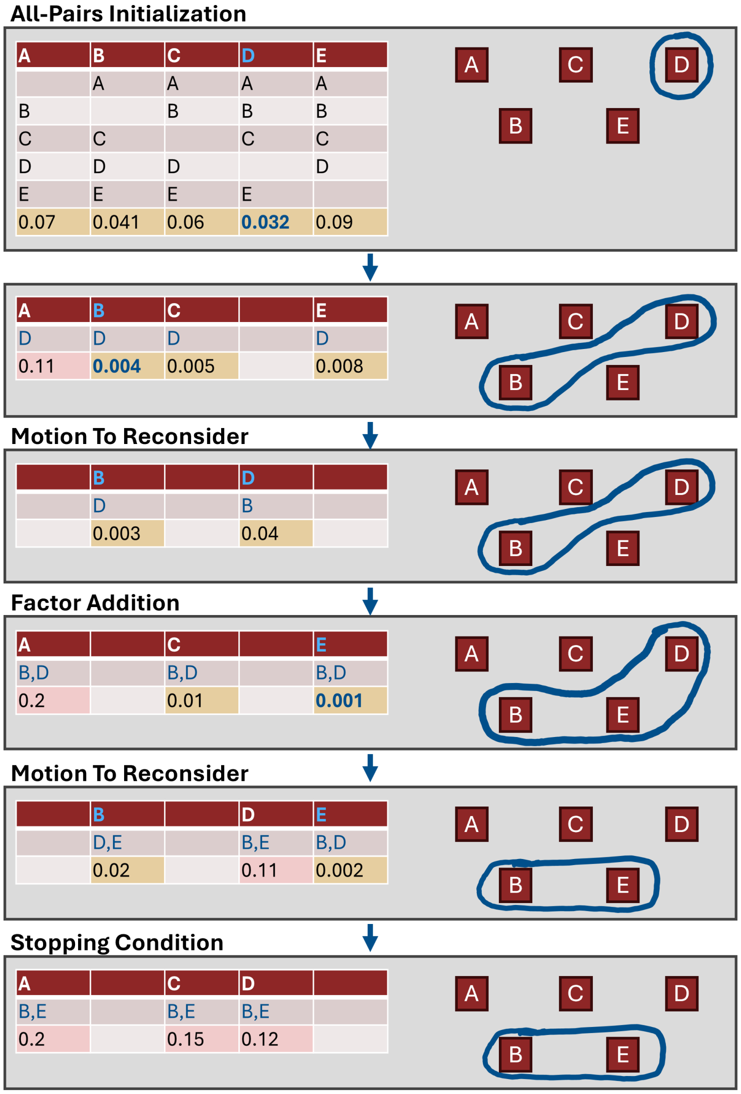


**Supplementary Figure S1. Factor Selection Procedure.** This figure shows an example of the Factor Selection Procedure with five potential factors. The table on the left shows the tests that were conducted, with the top (dark red) row showing which factors are being tested. The cells below show the factors that are included in the test but are not being tested. The bottom-most row is the average p-value of all the tests in that column (apart from the all-pairs initialization, this average is only of one test). The boxes on the right represent the factors, and the blue line encloses the factors in the hypothesis at the end of those tests. This graphic demonstrates the ways that the hypothesis changes during the hypothesis-building procedure.

***A.3. Estimation*** When the parameters are estimated with the Estimation Group, each time window is assigned the aggregate set of factors and allowed to estimate freely with those factors. Then, that estimation is run twice more, with the initial conditions of the new estimation being the results of the previous estimation. The second time, the number of subphases for estimation was reduced from $4$ to $1$, the size of the first subphase of the estimation phase was increased from $2.52 \times (p+7)$ to $5.04 \times (p+7)$ ($p$ is the number of parameters being estimated), and the number of iterations for estimating the standard error was increased from $1000$ to $5000$. This is known as the convergence reduction and is standard practice in the Rsiena literature [13]. This procedure was conducted three times for each time window and the estimation with the lowest convergence was selected as the estimate.

### Supplementary Note S3: Procedure Validation

**A Simulated Dataset** The validation of the Factor Selection Procedure was conducted by producing a set of simulated time windows where the underlying set of factors is known. To create these data, a time point 1 is selected at random from the pool of real networks from the dataset. Then, a set of factors and factor weights is selected. Using this set of factors and factor weights, a simulated time point 2 is produced. These simulated data were produced in group model sets where multiple time point 2s were produced from different time point 1s with the same underlying model.

Our simulated dataset consists of 185 group model sets with at least 20, but up to 50, examples of each model. There are many points in the process where the computation can fail, and these dataset numbers exclude the examples that did not make it

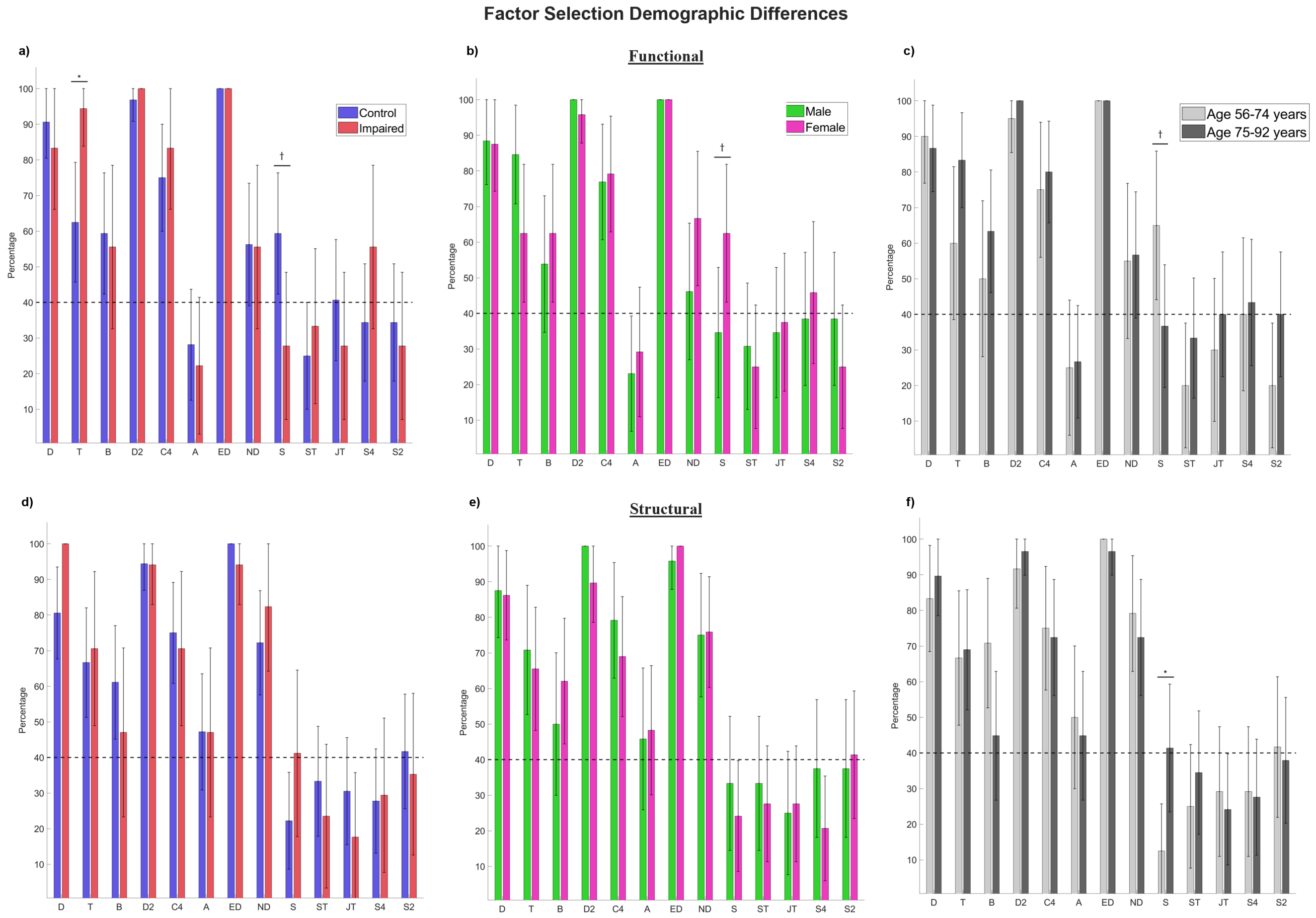


**Supplementary Figure S2. Factor Inclusion Comparisons.** These are the results of the analysis of the inclusion and exclusion of factors based on the demographic measures of impairment group (Figures a and d), sex (Figures b and e), and age group (Figures c and f). Figures (a-c) are for the functional networks, and Figures (d-f) are the structural networks. Five analyses showed significant differences or trends between groups —prevalence of Transitive Triads between impairment ($p = 0.033$) and prevalence of Same Subnetwork between impairment ($p = 0.064$), sex ($p = 0.091$), and age groups ($p = 0.094$) in the functional, and prevalence of Same Subnetwork between age groups ($p = 0.044$) in the structural.

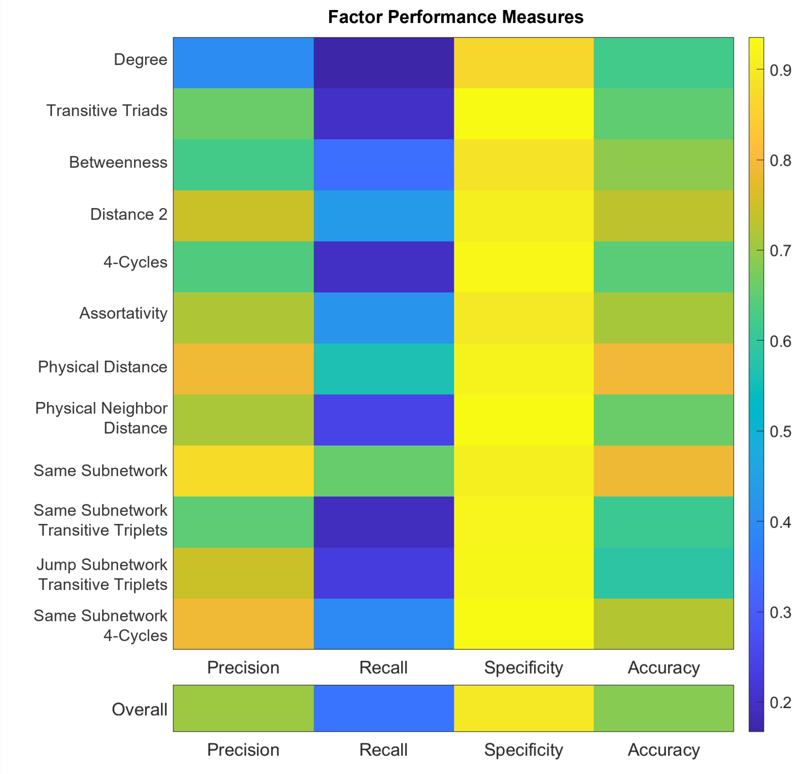


**Supplementary Figure S3. Overall factor performance.** This graph shows the relative success on metrics of precision, recall, specificity, and accuracy for individual factors. The lowest separated row represents the overall success of all the individual examples.

through the procedure.

Factor weights for the simulated data were selected based on a baseline heuristic of the behavior of the factor when it is run independently in order to accommodate the scale differences between factors. The rates for the simulated data are $\rho = 5$, $\rho = 20$, $\rho = 35$, $\rho = 50$, and $\rho = 65$ to cover a wide range of change behavior. Each time a model was selected to simulate, a desired number of factors was specified, and that many factors were then randomly selected. Thus, we controlled the number of factors that the models had, but we did not control the factors that would be selected. The dataset was designed to contain models with various numbers of factors.

Certain factors are underrepresented in the dataset—Factors Euclidean Distance ($ED$), Euclidean Neighbor Distance ($ND$), and Same Subnetwork 4-Cycles ($S4$) have fewer simulated examples, as they were added to the analysis later in the process. The simulated dataset also contains more functional networks at time point 1s than structural networks, as these were also included later. This simulated dataset is not a direct analog to the real dataset; however, it serves as a reasonable baseline for understanding the factor selection procedure and informing future steps in analysis.

**B Simulated Evaluation** We used measures of *precision*, *recall*, *specificity*, and *accuracy* to evaluate the success of the procedure. Figure S3 shows the relative performance of each factor on these metrics. It can be seen that all factors perform poorly on recall and very well on specificity, evidence that the Factor Selection Procedure often misses factors that should be included but rarely includes factors that should be excluded.

These results only provide a surface-level analysis of the overall performance of the procedure, with excellent specificity and better precision than recall. However, a threshold value on the percentage of individual simulated time windows in a group model set was used to develop the receiver operating characteristic (ROC) and precision-recall curves for the group model sets (Figure S4). The ROC curve and the precision-recall curve are commonly used tools for determining the success of a particular algorithm or procedure [14–16].

The area under these curves is considered a demonstration of the procedure's performance, with $AUC = 0.5$ being a procedure of no skill and $AUC = 1$ being a procedure of perfect skill [17, 18]. As stated, the Factor Selection Procedure has an area under the ROC curve of 0.767 and an area under the precision-recall curve of 0.787.

The F-score ($F_\beta$) calculates the performance of the procedure on various scales of precision and recall.

$$F_\beta = \left(1+\beta^2\right) \frac{PR \times RE}{\beta^2 \times PR + RE} \tag{S1}$$

where $\beta$ represents the fractional priority of recall over precision. A $\beta = 1$ represents the success of the procedure with equal

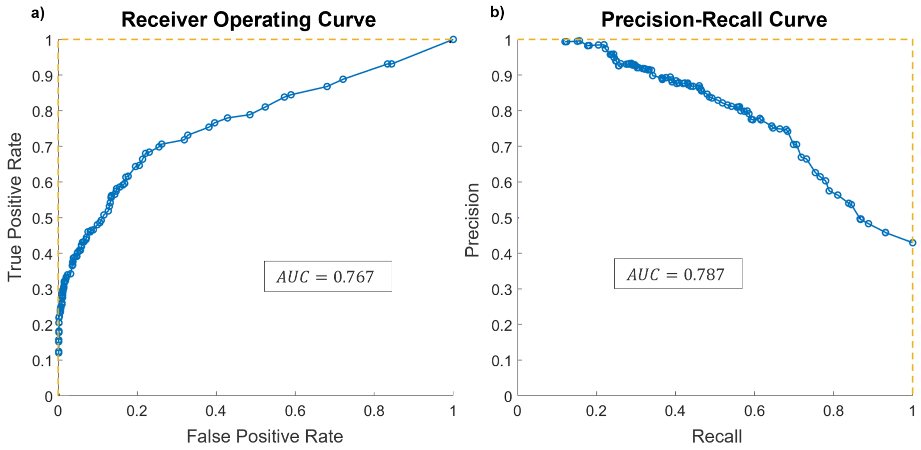


**Supplementary Figure S4. Simulated performance.** Receiver Operating Characteristic (ROC) and Precision-Recall Curve for the simulated data to validate the Factor Selection Procedure. ROC area under the curve: $AUC = 0.767$; PR area under the curve: $AUC = 0.787$.

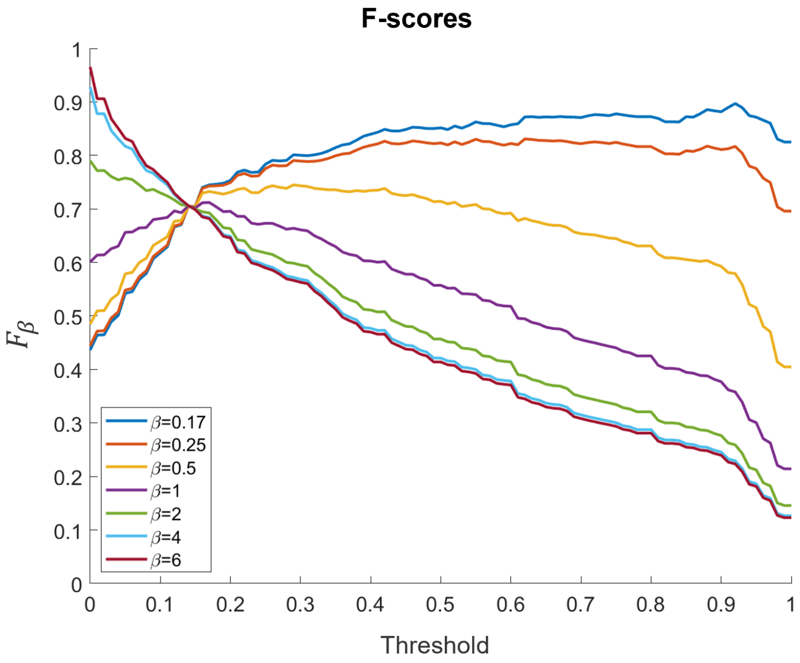


**Supplementary Figure S5. Overall F-score.** The F-score for the overall factor selection procedure at various values of $\beta$ demonstrates the superior performance of precision over recall and the differences in overall performance at different thresholds.

priority of recall and precision. A $\beta > 1$ represents the success of the procedure if recall is considered more important than precision. A $\beta < 1$ represents the success of the procedure if precision is considered more important than recall [15].

The F-score may be calculated at any threshold value, and Figure S5 shows $F_\beta$ graphed as a function of threshold. Various values of $\beta$ are included to demonstrate the performance based on different priorities of precision and recall. It is evident from the graph that prioritizing precision over recall gives a higher overall performance at most thresholds. This is consistent with the observation that the procedure generally performs better at precision than recall.

Because of the interdependence of the various factors, it is possible that the influence of a particular factor may be partially or mostly explained by another factor, thus increasing the likelihood that one of those factors will be missed. This indicates that some models in the simulated dataset are less derivable than others, but the interaction between factors suggests that even a derived model that is incorrect by missing factors can still explain most of the changes occurring on the network. This is in line with the factor selection procedure's criteria defined, as it is, as only including the minimum set of factors that can be considered not to be zero.

A threshold of $40\%$ is applied after the factor selection procedure such that a factor is included in the aggregate hypothesis if $40\%$ of time windows contain a particular factor. This $40\%$ threshold was selected based on these simulated results, focusing on the F-score with $\beta = 0.5$, prioritizing precision over recall (Figure S5).

**Supplementary Table S1. Factor Equations.** The equations for the different factors found in the brain data. All factors were found for both the functional and the structural except Assortativity, which was only found in the structural and Same Subnetwork and Same Subnetwork 4-Cycles which were only found in the functional. The equations were adapted from the RSiena manual [13] and represent the calculated value in the objective function for the selected node $i$. $\boldsymbol{G}_{ij}$ is the value for the connection between node $i$ and node $j$, $\boldsymbol{X}_{ij}$ is the Euclidean distance between node $i$ and node $j$, and $v_i$ is the subnetwork of node $i$ with $[v_i = v_j]$ indicating the node $i$ and node $j$ are in the same subnetwork.

| Factor | Equation | Weight Sign |
|---|---|---|
| Degree | $s_i = \sum_{j=1}^{N} \boldsymbol{G}_{ij}$ | Negative |
| Transitive Triads | $s_i = \sum_{j=1}^{N} \sum_{h=j+1}^{N} \boldsymbol{G}_{ij}\boldsymbol{G}_{ih}\boldsymbol{G}_{jh}$ | Positive |
| Betweenness | $s_i = \sum_{j=1}^{N}\sum_{h=1}^{N} \boldsymbol{G}_{hi}\boldsymbol{G}_{ij}\left(1-\boldsymbol{G}_{hj}\right)$ | Negative |
| Distance 2 | $s_i = \sum_{j=1}^{N} (1-\boldsymbol{G}_{ij}) \max_h \left(\boldsymbol{G}_{ih}\boldsymbol{G}_{hj}\right)$ | Negative |
| 4-Cycles | $s_i = \frac{1}{4}\sum_{j=1}^{N}\sum_{k=1}^{N}\sum_{h=1}^{N} \boldsymbol{G}_{ij}\boldsymbol{G}_{ik}\boldsymbol{G}_{hj}\boldsymbol{G}_{hk}$ | Negative |
| Assortativity | $s_i = \sum_{j=1}^{N} \boldsymbol{G}_{ij} \left(\sum_{h=1}^{N} \boldsymbol{G}_{ih}\right)^{1/2} \left(\sum_{k=1}^{N} \boldsymbol{G}_{jk}\right)^{1/2}$ | Negative |
| Euclidean Distance | $s_i = \sum_{j=1}^{N} \boldsymbol{G}_{ij}(\boldsymbol{X}_{ij} - \sum_{h=1}^{N} \boldsymbol{X}_{ih})$ | Negative |
| Euclidean Neighbor Distance | $s_i = \sum_{j=1}^{N}\sum_{h=1}^{N} \boldsymbol{G}_{ij}\boldsymbol{G}_{ih}\boldsymbol{X}_{hj}$ | Positive |
| Same Subnetwork | $s_i = \sum_{j=1}^{N} \boldsymbol{G}_{ij}\left[v_i = v_j\right]$ | Positive |
| Same Subnetwork 4-Cycles | $s_i = \frac{1}{4}\sum_{j=1}^{N}\sum_{k=1}^{N}\sum_{h=1}^{N} \boldsymbol{G}_{ij}\boldsymbol{G}_{ik}\boldsymbol{G}_{hj}\boldsymbol{G}_{hk}\left[v_i = v_h\right]$ | Negative |

A simulated dataset allows this procedure to be evaluated for its precision, recall, specificity, and accuracy when the underlying model is known. The procedure is currently specific to undirected networks, and the evaluation of the simulated data was conducted with the goals and assumptions of studying brain networks. However, the basics of this procedure may be useful in other applications with other kinds of networks. It is a proof of concept for applying the SAOM without a specific hypothesis in mind and having the development and derivation of a specific hypothesis be data-driven. Overall, the factor selection procedure is found to be a useful and effective tool for applying the SAOM to brain networks.

### Supplementary Note S4: Factor Equations

The factor equations for the factors found 40% of the Factor Group are included in Table S1. These equations were adapted from the Rsiena manual [13] but apply only for the undirected networks.
